# Mechanism of nucleus-chloroplast communication by alternative promoter usage and stromules to establish photomorphogenesis in Arabidopsis

**DOI:** 10.1101/2024.05.13.593997

**Authors:** Jae-Hyung Lee, Thu Minh Doan, Sandhya Senthilkumar, Chan Yul Yoo

## Abstract

Interorganellar communication is essential for maintaining cellular and organellar functions and adapting to dynamic environmental changes in eukaryotic cells. In plants, light triggers photomorphogenic development, including chloroplast biogenesis and the inhibition of hypocotyl elongation, through photoreceptors such as the red/far-red-sensing phytochromes and their downstream signaling pathways. However, the mechanism of interorganellar crosstalk underlying photomorphogenesis remains elusive. Here, we investigate the role of light-regulated alternative promoter usage in *NUCLEAR CONTROL OF PEP ACTIVITY* (*NCP*), a gene encoding a phytochrome signaling component that is dual-localized to the nucleus and chloroplasts. The long transcript variant (*NCP-L*) is upregulated under high red light, while the short variant (*NCP-S*) predominates in dark or low red light conditions. This light-regulated alternative transcription initiation of *NCP* is dependent on PHYTOCHROME-INTERACTING FACTORS (PIFs). The NCP-L isoform primarily localizes to chloroplasts, whereas the NCP-S isoform is found in the cytoplasm and nucleus. Notably, chloroplast-localized NCP-L translocates to the nucleus via stromules. Consequently, NCP-L, present in both chloroplasts and the nucleus, initiates chloroplast biogenesis and inhibits hypocotyl growth during photomorphogenesis, whereas NCP-S is nonfunctional and degraded by the 26S proteasome. Taken together, our findings elucidate the mechanisms by which light-regulated *NCP* alternative promoter usage and NCP retrotranslocation via stromules control photomorphogenesis in Arabidopsis. These mechanisms provide insights into interorganellar communication, orchestrating organ-specific developmental processes in response to fluctuating light environments.

## Introduction

The eukaryotic cell is distinguished by membrane-bound organelles, including the nucleus, endoplasmic reticulum, vacuoles, Golgi apparatus, and mitochondria. In plants and algae, chloroplasts are essential organelles that evolved from a free-living cyanobacterial ancestor through endosymbiosis and are responsible for photosynthesis (1). Chloroplasts are semi-autonomous, as most genes in the plastidial genome have been either lost or transferred to the nuclear genome during the co-evolution of plastids and their host cells (2). However, the plastidial genome still retains genes crucial for chloroplast biogenesis and function. This dual genetic control necessitates coordinated gene expression from both the nuclear and plastidial genomes to regulate chloroplast biogenesis, function, and homeostasis (3). Furthermore, chloroplasts communicate bidirectionally with the nucleus in response to internal and external stimuli to regulate nuclear gene expression (4). This anterograde and retrograde interorganellar communication is essential for orchestrating plant growth and development, as well as facilitating adaptation to environmental changes (5).

Light is an essential environmental cue for plants and initiates developmental programs involving chloroplast biogenesis (6). When seeds germinate under the ground (dark), a developmental program called skotomorphogenesis promotes hypocotyl elongation to allow seedlings to emerge from the soil, and it inhibits chloroplast biogenesis to prevent photooxidative damage when seedlings encounter light later. When seedlings are exposed to light, a developmental program called photomorphogenesis is activated to restrict hypocotyl growth and initiate the differentiation of plastids into photosynthetically active chloroplasts (7). Hypocotyl growth is controlled by genome-wide transcriptional changes in the nucleus (8). Chloroplast biogenesis requires the synthesis of plastid-localized proteins to generate functional photosynthetic components such as the light-harvesting chlorophyll a/b complex, photosystems, ribulose bisphosphate carboxylase/oxygenase, and thylakoid membranes where photosynthetic machineries are built. These genes encoding photosynthetic components are transcribed from the nuclear and plastidial genomes, which are called photosynthesis-associated nuclear genes (PhANGs) and plastid genes (PhAPGs), respectively (3, 9). Establishing photomorphogenesis requires coordinated gene expression from the nuclear and plastidial genomes (9).

Light is perceived by a suite of photoreceptors, including the red- and far-red-light-sensing phytochromes (PHYs) (10). While the inactive form of PHYs in the dark is localized in the cytoplasm, photoactivated PHYs translocate into the nucleus and form condensates called photobodies (10, 11). PHY signaling is activated to reprogram the nuclear genome by regulating the stability and activity of PHYTOCHROME INTERACTING FACTORS (PIFs), which are master repressors of photomorphogenesis (12, 13). In the dark, most PIFs are abundant and activate growth-related genes in the nucleus to promote hypocotyl growth. PIFs are also known to repress both nuclear and plastidial genes associated with photosynthesis and chloroplast biogenesis. Thus, photoactivated PHYs repress the stability and activity of PIFs in the nucleus, triggering nucleus-to-plastid anterograde signaling to activate plastid gene expression (9).

Plastid genes are transcribed by the plastid-encoded RNA polymerase (PEP) and the nuclear-encoded RNA polymerase (NEP) (14). Recently, the structure of the PEP complex in plants has been reported in cryoelectron microscopy (cryo-EM) studies in tobacco and white mustard (15–17). The PEP originated in bacteria and is a multisubunit RNA polymerase complex consisting of 4 bacterial-like core subunits and plant-specific PEP-associated proteins (PAPs). The bacterial-like core genes (*rpoA*, *rpoB*, *rpoC1*, and *rpoC2*) are plastid genes transcribed by the other, phage-type RNA polymerase, the NEP. The PAPs, encoded in the nuclear genome, are imported into chloroplasts and assembled with the core proteins to yield a functional PEP holoenzyme complex. PAPs are considered structural components required for the assembly and stability of the PEP complex and other enzyme activities (15–17). PEP assembly is triggered by light via PHY signaling and is repressed by dark via PIFs in the nucleus (18). These characteristics suggest that nucleus-chloroplast interorganellar communication is crucial to establish photomorphogenesis.

A forward genetic screen to identify PHY signaling components for chloroplasts previously identified REGULATOR OF CHLOROPLAST BIOGENESIS (RCB) and NUCLEAR CONTROL OF PEP ACTIVITY (NCP) in Arabidopsis (18, 19). Interestingly, RCB and NCP are dual-localized to the nucleus and plastids, where they play a role in two major photomorphogenic developmental processes: hypocotyl growth inhibition and chloroplast biogenesis. In the nucleus, both RCB and NCP are required for the formation of large phyB photobodies and the degradation of PIF1 and PIF3 to inhibit hypocotyl elongation. PHY-mediated PIF degradation in the nucleus sends nucleus-to-plastid signals to active *PhAPGs* expression by triggering PEP assembly and independently by activating sigma factors for promoter recognition (18, 20). RCB activates PEP assembly primarily in the nucleus by degrading PIFs (18). Notably, NCP plays pivotal dual roles in controlling PIF degradation in the nucleus and PEP assembly in plastids (19). These genetic studies laid the foundational framework of nucleus-chloroplast communication (9). However, the mechanism of nucleus-to-plastid signaling remains to be understood. The dual localization of these proteins raises the question of how light regulates the nuclear and chloroplast localization of these PHY signaling components to coordinate hypocotyl growth inhibition in the nucleus and chloroplast biogenesis in the plastids.

In plants, PHYs induce genome-wide changes in light-dependent alternative splicing and alternative promoter selection, which generates distinct transcript variants and thus protein isoforms with diverse functions and subcellular localizations (21, 22). This observation prompted us to test whether the dual localization of RCB and NCP is regulated at the transcript level via alternative splicing or alternative promoter usage. We found alternative transcription start sites in *NCP*, but not in *RCB*; in *NCP*, they produce a long NCP isoform with an N-terminal chloroplast transit peptide and a short isoform without the N-terminus. Surprisingly, the long NCP isoform primarily localizes to chloroplasts and then translocates to the nucleus via stromules to initiate chloroplast biogenesis and inhibit hypocotyl growth in the nucleus. In contrast, the cytoplasm/nucleus-localized short NCP isoform is not functional and is degraded by the 26S proteasome. Therefore, our results demonstrate the dynamic regulation of interorganellar communication via alternative promoter usage and retrotranslocation through stromules. Our findings reveal the framework of the mutual signaling cascades between the nucleus and plastids to initiate chloroplast biogenesis and establish photomorphogenesis.

## RESULTS

### Light regulates the alternative transcription initiation of *NCP* in a PIF-dependent manner

The nucleus and chloroplast dual-localized NCP and RCB paralogs possess a chloroplast-targeting transit peptide (cTP) in their N termini and a putative nuclear localization signal (NLS) in the middle (18, 19). We hypothesized that different NCP or RCB isoforms with or without the cTP could be produced via PHY-mediated alternative promoter selection (21). We first tested whether *NCP* or *RCB* generate different transcript variants by using 5′ rapid amplification of cDNA ends (5′ RACE) analysis of total RNA extracted from seedlings grown under white light (WL) and darkness. While a single *NCP* transcript was detected in seedlings grown under WL, two *NCP* transcript variants were detected in the dark condition (Figure 1A). We cloned and sequenced the longer *NCP* cDNA (*NCP-L*) and the shorter *NCP* cDNA (*NCP-S*) to determine the difference between the two transcript variants. The *NCP-L* transcript started within the region for the transcription start sites (TSSs) annotated in TAIR (referred to here as TSS1) (Figure 1B). The *NCP-S* transcript started with TSS2 in the region after the first ATG in the coding sequence (Supplemental Figure 1A), which led to production of short isoform with a truncated N-terminus (Supplemental Figure 2). We also analyzed the *RCB* transcript under same conditions using 5’-RACE analysis and found only a single *RCB* transcript in both WL and dark conditions (Figure 1C). Interestingly, the sequencing of multiple clones from the 5’-RACE analysis detected the *RCB* TSS downstream of the annotated TSS (Supplemental Figure 1B). These results indicate that two *NCP* transcript variants with different TSSs and one *RCB* transcript exist in the dark, but not in the light.

**Figure 1.**
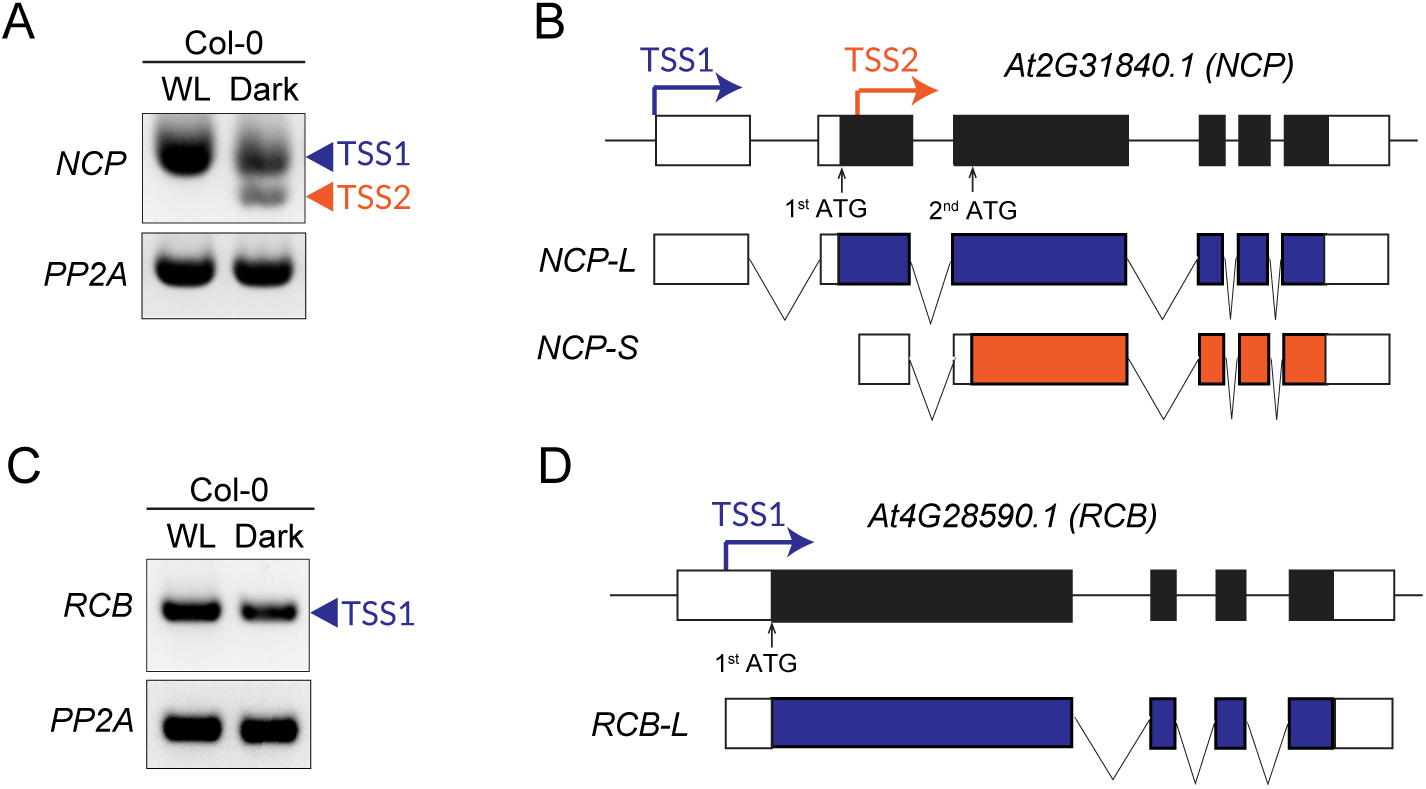
Identification of transcript variants of *NCP* and *RCB*. **(A)** Identification of *NCP* transcript variants with different transcription start sites. Wild-type (Col-0) plants were grown on MS-agar plates in either 100 μmol m^−2^ s^−1^ white light (WL) or darkness for 4 d before extracting total RNA from whole seedlings. Two specific bands were amplified by 5’ RACE-PCR. The specific bands were confirmed by Sanger sequencing, and the transcription start sites were designated as TSS1 (blue arrowhead) and TSS2 (red arrowhead). *Protein phosphatase 2A* (*PP2A*) is shown as an internal control. **(B)** Schematic representation of *NCP* transcript variants with different transcription start sites. The longer and shorter *NCP* transcripts represent *NCP-L* and *NCP-S*, respectively. Predicted start codons corresponding to the different transcription start sites are depicted below. Untranslated regions are indicated as white boxes, and constitutive exons are marked as dark boxes. Introns are shown as horizontal lines. **(C) and (D)** 5’ RACE analysis of the *RCB* gene. Seedling growth and RNA extraction were performed as described in **(A)**. The identified transcription start site was designated as TSS1 (blue arrowhead) **(C)**. *PP2A* was used as an internal control. The longer *RCB* transcript cloned by 5’ RACE is represented as “*RCB-L*” **(D)**.

We next tested whether the *NCP-L* transcript observed under WL could be due to higher light intensity or light quality (strong blue light in WL). We performed 5’ RACE analysis of seedlings grown under 70 µmol m^−2^ s^−1^ red light (R70), 20 µmol m^−2^ s^−1^ R light (R20), or dark conditions in three independent biological replicates and quantified the percentages of the two transcripts. The *NCP-L* transcript, which included TSS1, was expressed strongly under the R70 condition, decreased in the weaker R20 condition, and decreased even more in the dark condition (Figure 2A). In contrast, *NCP-S* was detected in the dark and in weaker R20 light but was not detected in the higher R70 light condition (Figure 2A). More than 60% of the transcripts produced in the dark were generated with TSS2, while more than 70% of the transcripts in R20 and 100% of transcripts in R70 were generated with TSS1 (Figure 2B). Our result is consistent with a previous report showing that TSS2 in *NCP* is strongly observed in wild-type grown in the dark and in *phyA-211/phyB-9* in red light (21).

**Figure 2.**
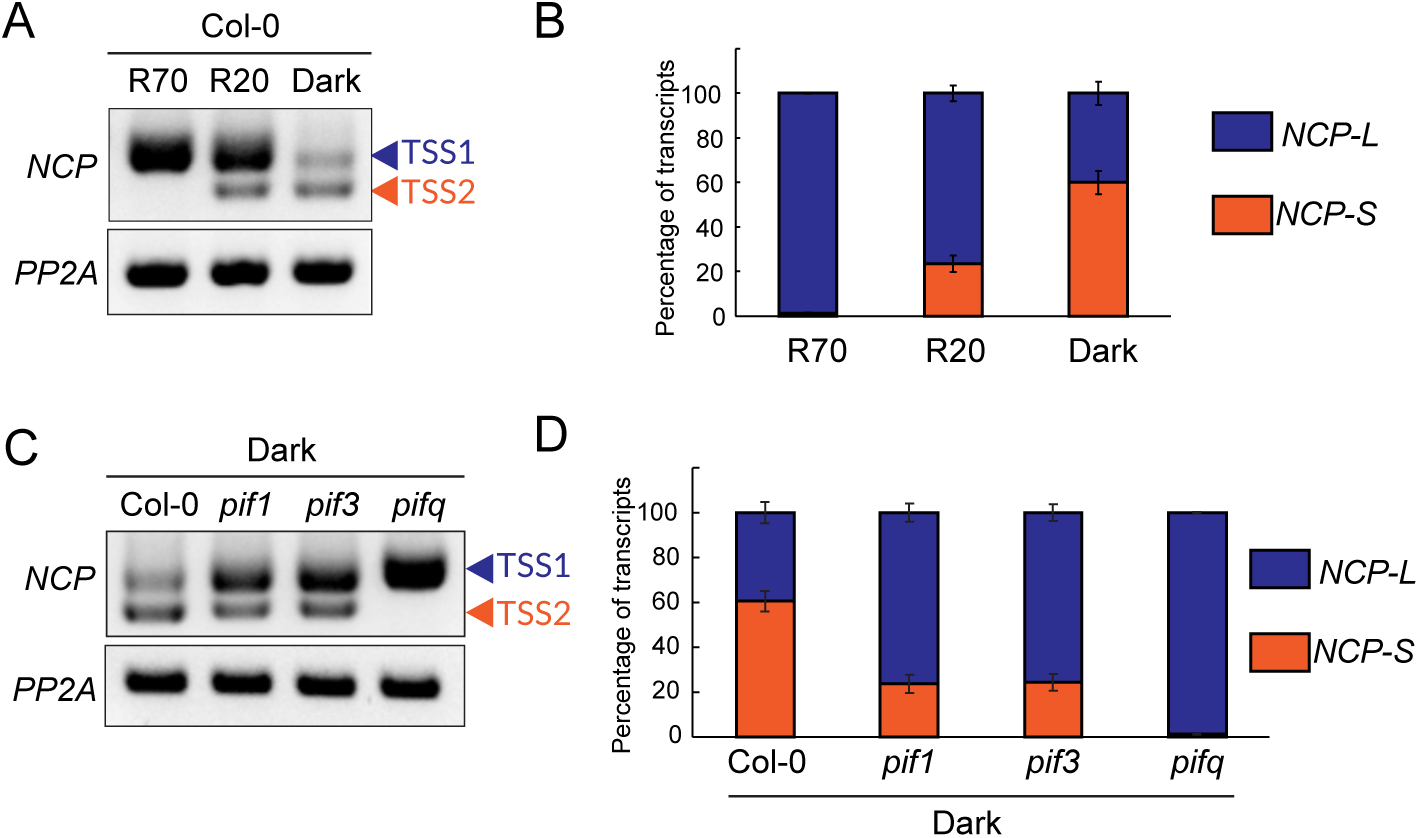
Light controls alternative transcription start sites in *NCP* via a PIF-dependent mechanism. **(A)** Effect of light intensity on the use of the alternative *NCP* promoter. Col-0 plants were grown on MS-agar plates in 70 μmol m^−2^ s^−1^ red light, 20 μmol m^−2^ s^−1^ red light, or true-dark conditions for 4 d before extracting total RNA from whole seedlings. The transcription start sites defined by 5’ RACE analysis were designated TSS1 (blue arrowhead) and TSS2 (red arrowhead). *PP2A* was used as an internal control. R, red light. **(B)** Quantification of longer and shorter *NCP* transcripts. Quantification was performed using 5’ RACE-PCR as described in **(A)**. DNA blots on the three 5’ RACE-PCR samples were quantified and averaged using ImageJ software. Blue and red bar graphs indicate the quantified band ratios for the longer (*NCP-L*) and shorter (*NCP-S*) *NCP* transcripts, respectively. Bars indicate the SD of three biological replicates. **(C)** Effects of *PIF* mutations on alternative promoter usage in *NCP*. Col-0, *pif1*, *pif3*, and *pifq* plants were grown on MS-agar plates for 4 d in darkness before extracting total RNA from whole seedlings. Each transcription start site identified by 5’ RACE-PCR is marked as TSS1 (blue arrowhead) or TSS2 (red arrowhead). *PP2A* was used as an internal control. **(D)** Quantification of longer and shorter *NCP* transcripts in *PIF*-defective mutants. Quantification was performed using the three independent 5’ RACE-PCR samples described in **(C)**. Blue and red bar graphs indicate the quantified band ratios for the longer and shorter *NCP* transcripts, respectively. Bars indicate the SD of three biological replicates.

Since the central mechanism of light signaling is to repress the levels and activities of a family of antagonizing transcription factors called PHYTOCHROME INTERACTING FACTORS (PIFs), we next asked whether PIFs repress the light-regulated alternative TSSs in *NCP* by using *pif1, pif3*, and *pifq* (*pif1 pif3 pif4 pif5*) mutants grown in the dark. The *NCP-L* transcript level increased marginally in the *pif1* and *pif3* single mutants, and it increased substantially in the *pifq* mutant (Figure 4C). In contrast, the *NCP-S* transcript completely disappeared in the *pifq* mutant (Figure 4C). The *NCP-S* transcript accounted for approximately 60% of all *NCP* transcripts in Col-0 under dark conditions; this level was reduced to approximately 25% in both the *pif1* and *pif3* mutants and completely disappeared in the *pifq* mutant (Figure 4D). Taken together, these results demonstrate that PHY-mediated red light signaling controls alternative transcription initiation of *NCP* in favor of TSS1 under higher light conditions and TSS2 under lower R light and dark conditions in a PIF-dependent manner.

### NCP isoforms localize to different subcellular compartments

NCP has been previously reported to be a dual-targeted protein (19), but the mechanism underlying its dual targeting remained unclear. The *NCP-L* transcript encodes an isoform containing both a cTP (amino acids 1-48) and a putative NLS (amino acids 118-145), whereas the *NCP-S* transcript encodes an isoform containing only the NLS because the 2^nd^ methionine (start codon) is located in the 56^th^ residue of the NCP-L coding sequence (Supplemental Figure 2). Thus, we hypothesized that light-regulated alternative transcription initiation would produce nucleus-localized NCP-S and chloroplast-localized NCP-L isoforms as a mechanism of dual targeting. We attempted to examine the subcellular localization of NCP-L and NCP-S by expressing these two proteins fused with cyan fluorescent protein (CFP) in tobacco leaves. The NCP-L-CFP signal was mostly detected in chloroplasts and often observed in the nucleus when it was surrounded by chloroplasts (Figure 3A and Supplemental Figure 3A). Interestingly, we observed NCP-L-CFP localization in stromules (stroma-filled tubular extensions from the chloroplasts) that were connected to the nucleus (Figure 3A and Supplemental Figure 3B). In contrast, the NCP-S-CFP isoform was detected in the cytoplasm and the nucleus, but not in the chloroplasts (Figure 3A). We cannot exclude the possibility that nuclear signal of the NCP-S isoform is driven by the diffusion of NCP-S-CFP protein into the nucleus. We also tested both isoforms in Arabidopsis mesophyll protoplasts isolated from leaves. Interestingly, NCP-L-CFP was localized only in chloroplasts in Arabidopsis protoplasts, while NCP-S-CFP was localized in the cytoplasm and the nucleus (Figure 3B). However, stromules were not observed in our mesophyll protoplasts. This observation is consistent with previous reports showing that stromules are generally not induced in protoplasts without exogenous treatment (23). This result indicates that the primary location of the NCP-L-CFP is the chloroplasts due to the N-terminal cTP. Taken together, our results demonstrate that the NCP-L isoform localizes to chloroplasts first and then translocates to the nucleus via stromules, while the NCP-S isoform is localized in the cytoplasm and the nucleus.

**Figure 3.**
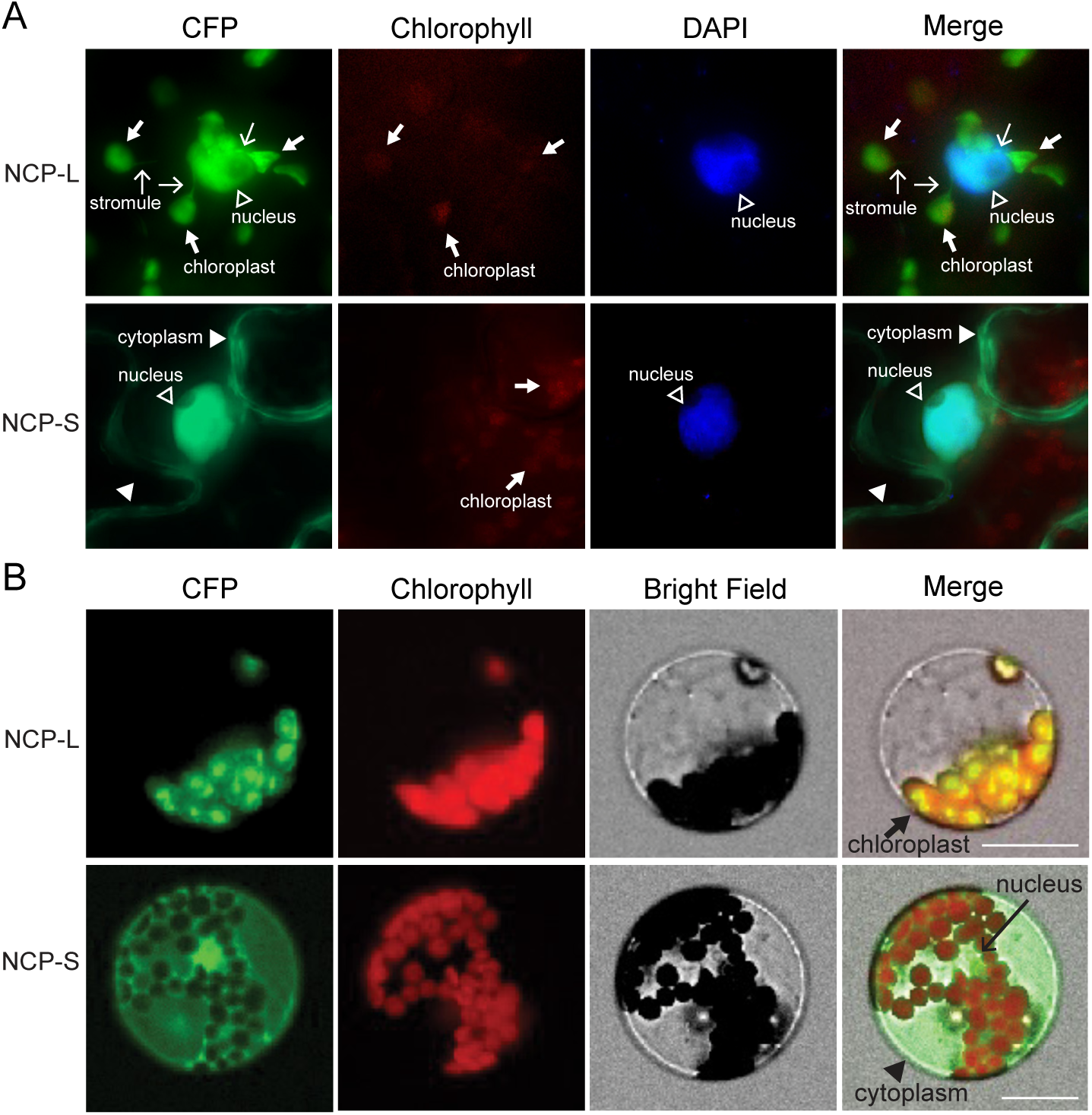
NCP-L and NCP-S protein isoforms show different subcellular localizations. **(A)** Distinct localizations of the NCP-L and NCP-S protein isoforms. The *UBQ10*-*NCP*-*L*-*CFP-FLAG* and *UBQ10*-*NCP*-*S*-*CFP-FLAG* fusion constructs were expressed transiently in tobacco leaves. Fluorescence microscopy images of epidermal cells were visualized. DAPI was used for staining nuclei. Thick arrows indicate chloroplasts. Thin arrows denote stromules. Empty arrowhead indicates nuclei. Filled arrowheads indicate a cytoplasmic signal. Chlorophyll autofluorescence is also shown. **(B)** Subcellular localization of the NCP-L and NCP-S isoforms in Arabidopsis mesophyll protoplasts. The same constructs used in **(A)** were expressed transiently in Arabidopsis protoplasts and visualized using fluorescence microscopy. Thick arrow indicates chloroplasts. Filled arrowhead denotes a cytoplasmic signal. Thin arrow indicates the CFP signal from the nuclei. Scale bars, 20 μm.

### The NCP-L isoform rescues both the chloroplast and nuclear phenotypes of the *ncp-10* mutant

The light-induced chloroplast/nuclear localization of the NCP-L isoform and the dark-favored cytoplasmic/nuclear localization of the NCP-S isoform led us to hypothesize that NCP-L could rescue the albino and tall hypocotyl phenotypes of the *ncp-10* null mutant, while NCP-S could rescue only its tall hypocotyl phenotype. To test our hypothesis on the chloroplast function of the NCP-L isoform, we generated transgenic plants expressing the ∼4.9-kb *NCP* genomic fragment fused to an HA-His tag with the 2^nd^ ATG mutated (ATG to TTG) to prevent the translation of the NCP-S isoform (*NCPpro:NCP*_2nd ATGm_*-HA-His*/*ncp-10*). As a control, we used transgenic plants expressing the wild-type *NCP* genomic fragment (*NCPpro:NCP-HA-His*/*ncp-10*), which fully rescued the albino phenotype and the defects in the PEP-dependent *psbA* and *rbcL* gene expression in *ncp-10* (Figure 4A and 4B). Transgenic plants expressing the mutated 2^nd^ ATG in the *NCP* genomic fragment also fully rescued the albino and PEP-dependent gene expression phenotypes of *ncp-10* (Figure 4A and 4B). These results indicate that the NCP-L isoform is responsible for chloroplast biogenesis.

**Figure 4.**
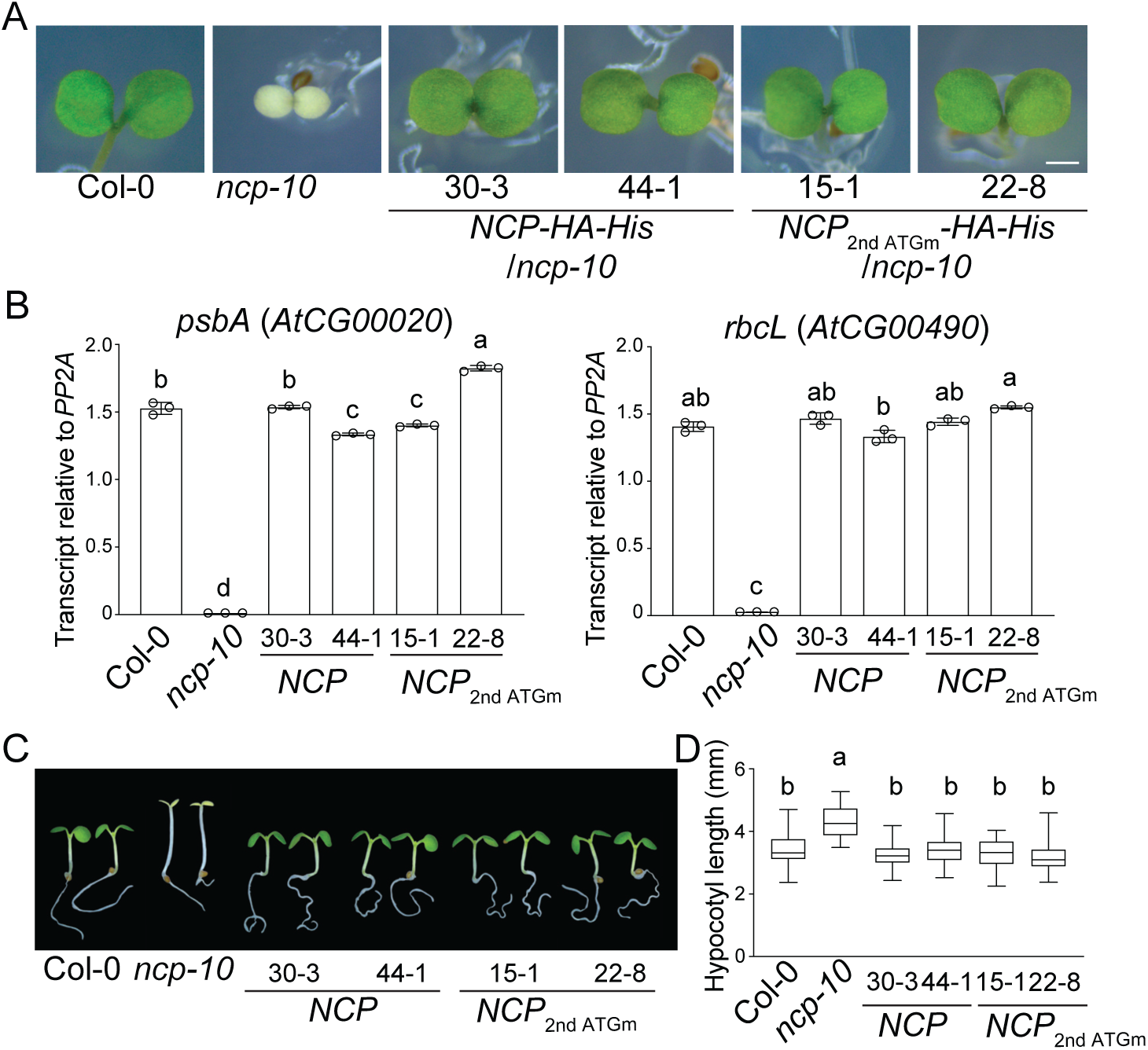
The NCP-L isoform is responsible for chloroplast biogenesis and hypocotyl elongation. **(A)** The *ncp-10* albino phenotype is rescued by expressing the wild-type NCP-L isoform (*NCPpro:NCP*-*HA*-*His*) and NCP-L with the 2^nd^ ATG mutated (*NCP*_2nd ATGm_-*HA*-*His*). Col-0, *ncp-10*, *NCPpro:NCP*-*HA*-*His*/*ncp-10*, and *NCPpro:NCP*_2nd ATGm_-*HA*-*His/ncp-10* seedlings were grown under 20 μmol m^−2^ s^−1^ R light for 4 d. Scale bar, 1 mm. **(B)** Transcript levels of PEP-dependent genes in *NCP*_2nd ATG-mut_-*HA*-*His ncp-10* plants. Seedlings were grown in 20 μmol m^−2^ s^−1^ R light for 4 d before harvesting whole seedlings for RNA extraction. Transcript levels were examined using qRT-PCR. Different letters denote statistically significant differences in the transcript levels (ANOVA, Tukey’s HSD, *P* ≤ 0.001). Error bars represent the SD of three biological replicates. **(C)** The *ncp-10* tall hypocotyl phenotype is rescued by expressing the wild-type NCP-L isoform (*NCPpro:NCP*-*HA*-*His*) and NCP-L with the 2^nd^ ATG mutated (*NCP*_2nd ATGm_-*HA*-*His*). Col-0, *ncp-10*, *NCPpro:NCP*-*HA*-*His*/*ncp-10*, and *NCPpro:NCP*_2nd ATGm_-*HA*-*His/ncp-10* seedlings were grown under 50 μmol m^−2^s^−1^ R light for 4 d. **(D)** Box-and-whisker plots showing hypocotyl measurements of the seedlings in (C). Different letters represent significant differences (*P* ≤ 0.001) determined by one-way ANOVA with post-hoc Tukey’s HSD test. Bars indicate SD (*n* = 30).

The tall hypocotyl of *ncp-10* under R light is attributable to NCP’s nuclear function in PHY signaling. Thus, we hypothesized that *NCP*_2nd ATGm_ transgenic plants would not fully rescue the tall hypocotyl of *ncp-10* due to the lack of translation of the cytoplasm/nucleus-localized NCP-S isoform. Surprisingly, the long hypocotyl phenotype of *ncp-10* was fully rescued both in wild-type and *NCP_2nd ATGm_* transgenic plants in the R light condition (Figures 4C and 4D). Since chloroplast-targeted NCP-L can translocate to the nucleus via stromules, these results suggest that the nuclear- and-chloroplast localization of the NCP-L isoform is sufficient for rescuing the tall-and-albino phenotypes of the *ncp-10* mutant. However, these data prompted us to question the biological function of the NCP-S isoform.

### Mistargeted NCP-S protein is nonfunctional and degraded via the 26S proteasome-dependent pathway

To determine the biological function of the NCP-S isoform, we generated transgenic plants expressing a *UBQ10*-promoter-driven *NCP-S* coding sequence fused with an HA-His tag in the *ncp-10* mutant background. As a control to compare the phenotypic contributions of NCP-L and NCP-S, we also generated transgenic plants expressing NCP-L in *ncp-10*. Consistent with *NCP_2nd ATGm_* transgenic plants, the transgenic plants expressing the NCP-L isoform showed fully rescued albino and tall hypocotyl *ncp-10* phenotypes (Figures 5A and 5B). However, the transgenic plants expressing the NCP-S isoform failed to rescue the albino phenotype (Figure 5A) or the tall hypocotyl phenotype of *ncp-10* under R light (Figures 5B and 5C). To determine whether this phenotypic difference was associated with the transcription or translation of the transgenes, we first analyzed the transcript levels of the transgenes using primers specific to *NCP-HA-His*. The transcript levels of the *NCP-L* and *NCP-S* transgenes were similarly overexpressed by the *UBQ10* promoters in both the *NCP-L* and *NCP-S* transgenic lines (Figure 5D). Surprisingly, however, the NCP-S-HA-His protein levels were much lower in the *NCP-S* transgenic lines than the NCP-L-HA-His protein levels in the *NCP-L* transgenic lines (Figure 5E). We thus asked whether the cytoplasm/nucleus-localized NCP-S protein could be degraded via the 26S proteasome pathway. We found that NCP-S protein accumulated with MG132, a 26S proteasome inhibitor, while the NCP-L protein level did not change (Figure 5E). Taken together, our results indicate that NCP-L localizes to chloroplasts first and then mature form of NCP (NCPm) after cleave of its cTP translocates to the nucleus to promote chloroplast biogenesis and then regulates hypocotyl growth inhibition via phytochrome signaling in the nucleus, while mistargeted NCP-S in the cytoplasm/nucleus is nonfunctional and degraded via the 26S proteasome-dependent pathway.

**Figure 5.**
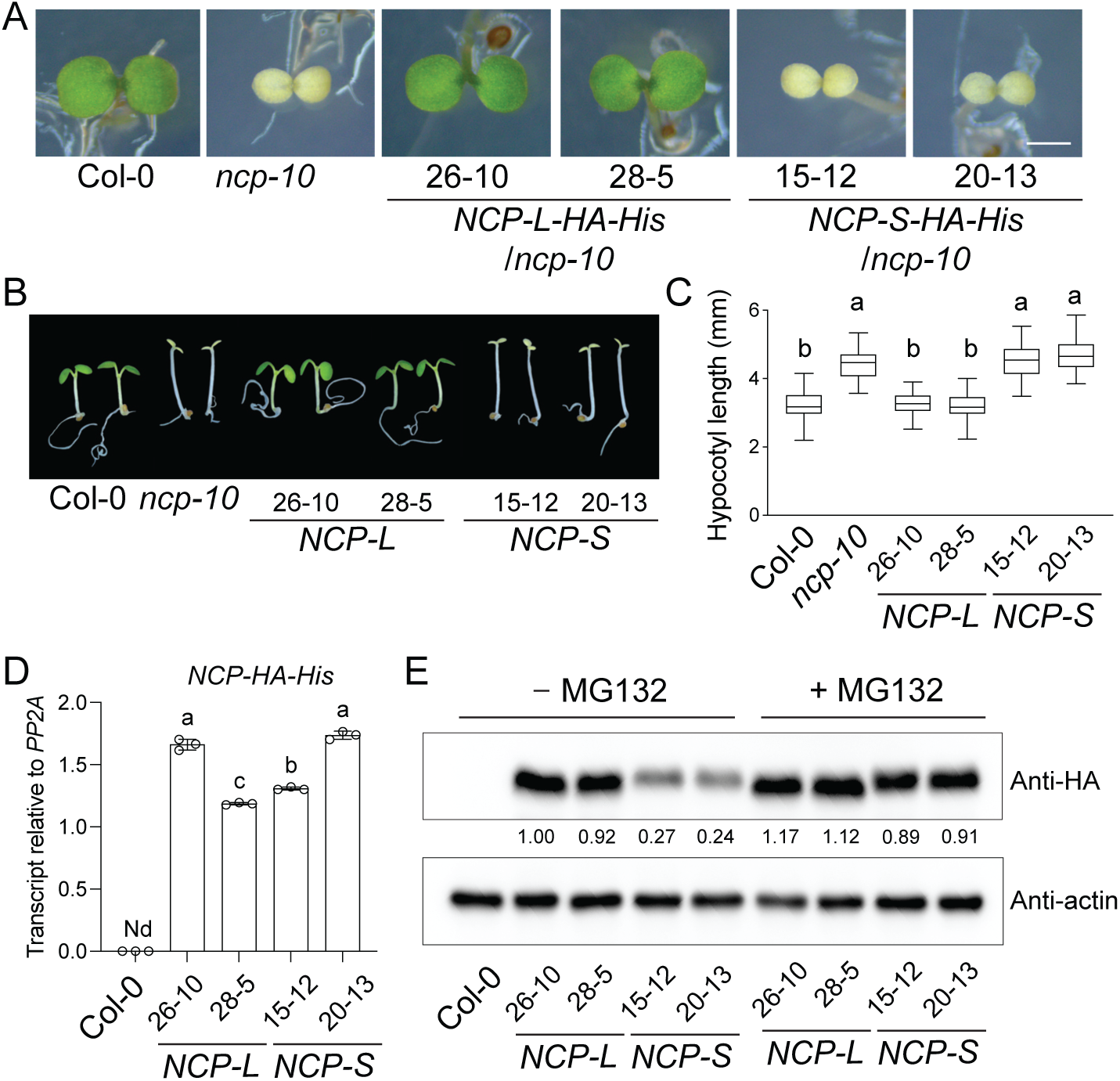
The mistargeted NCP-S protein isoform is nonfunctional and degraded via the 26S proteasome-dependent pathway. **(A)** Rescuing the *ncp-10* albino phenotype when overexpressing *UBQ10*-*NCP*-*L*-*HA*-*His*. Col-0, *ncp-10*, *NCP-L*-*HA*-*His/ncp-10*, and *NCP-S*-*HA*-*His/ncp-10* seedlings were grown under 20 μmol m^−2^ s^−1^ R light condition for 4 d. Scale bar, 1 mm. **(B)** Rescuing the *ncp-10* tall hypocotyl phenotype when overexpressing *UBQ10-NCP-L-HA-His*. Col-0, *ncp-10*, *NCP-L*-*HA*-*His/ncp-10*, and *NCP-S*-*HA*-*His/ncp-10* seedlings were grown under 50 μmol m^−2^s^−1^ R light for 4 d. **(C)** Box-and-whisker plots showing hypocotyl measurements of the seedlings in (B). Different letters represent significant differences (*P* ≤ 0.001) determined by one-way ANOVA with post-hoc Tukey’s HSD test. Bars indicate SD (*n* = 30). **(D)** Transcript levels of the *NCP*-*HA*-*His* transgene. Col-0, *UBQ10-NCP-L-HA-His,* and *UBQ10-NCP*-S*-HA-His* transgenic seedlings were grown under 50 μmol m^−2^ s^−1^ R light for 4 d and harvested for total RNA extraction. Transcript levels were examined using qRT-PCR. Different letters denote statistically significant differences in the transcript levels (ANOVA, Tukey’s HSD, *P* ≤ 0.001). Error bars represent the SD of three biological replicates. Nd, not detectable. **(E)** 26S proteasome-dependent degradation of the NCP-S isoform. Three-day-old Col-0, *UBQ10*-*NCP*-*L*-*HA*-*His,* and *UBQ10*-*NCP*-*S*-*HA*-*His* seedlings were incubated without and with 50 μM MG132 for 24 h before extracting total proteins from whole seedlings. Anti-HA antibody was used to detect NCP-L-HA-His and NCP-S-HA-His proteins. Actin was used as a loading control. Numbers indicate the relative protein level normalized to the actin level.

## Discussion

Interorganellar communication between the nucleus and chloroplasts is essential to establish photomorphogenesis, which involves hypocotyl growth regulation in the nucleus and chloroplast biogenesis from proplastids or etioplasts (5). In this study, we show that the nucleus-chloroplast dual-localization of NCP is regulated by light-controlled alternative promoter selection and retrotranslocation via stromules to establish photomorphogenesis in Arabidopsis. Light regulates alternative transcription initiation in *NCP* in favor of TSS1 via PIF-mediated phytochrome signaling, which produces the longer NCP-L isoform. NCP-L localizes to chloroplasts through the cTP and is converted to the mature form of NCP due to the cleavage of the cTP during chloroplast import. The mature form of NCP (NCPm) facilitates PEP assembly and activity to initiate chloroplast biogenesis. Then, NCPm translocates to the nucleus via stromules. In the nucleus, NCPm is required for PHY signaling to inhibit hypocotyl growth. In the dark, PHYs are inactivated, leading to PIF accumulation and hypocotyl elongation. PIFs inhibit TSS1 and promote *NCP* transcription from TSS2, producing a shorter NCP-S isoform that lacks an N-terminal cTP. Cytoplasmic NCP-S is nonfunctional and degraded via the 26S proteasome-dependent pathway. Together, these results outline the nucleus-chloroplast interorganellar communication mechanism that regulates NCP localization by light-regulated alternative promoter usage and retrotranslocation via stromules to establish photomorphogenesis (Figure 6).

**Figure 6.**
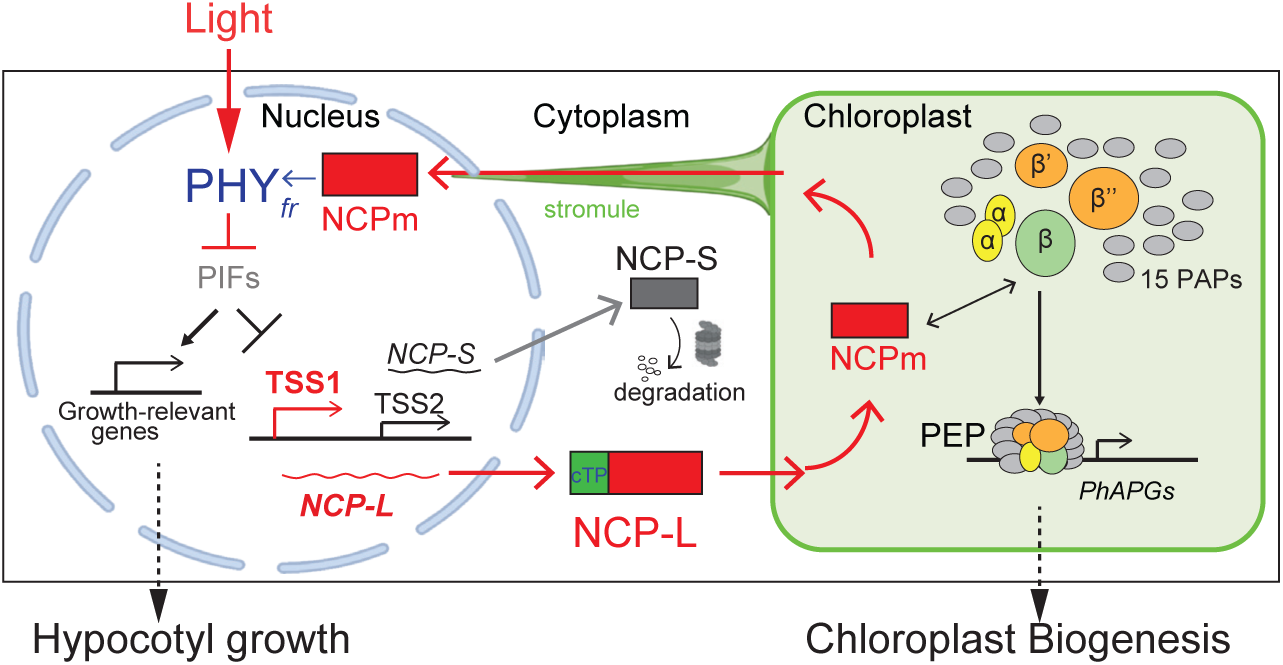
Model illustrating nucleus-chloroplast Interorganellar communication via alternative promoter usage and stromules during photomorphogenesis. Light triggers chloroplast differentiation by controlling PIF-dependent alternative promoter usage in *NCP*, producing chloroplast-targeted NCP-L. The mature form of NCP (NCPm) after cTP cleavage is required for PEP assembly to initiate chloroplast biogenesis. NCPm translocates to the nucleus via stromules and regulates PHY signaling to inhibit hypocotyl growth and establish photomorphogenesis.

Alternative promoter selection plays a crucial role in generating diverse transcripts that encode proteins with varied functions, subcellular localizations, and transcriptional/translational regulatory mechanisms (24–26). In plants, PHYs have been known to control alternative promoter selection to modulate protein localization as part of the adaptation to the light environment (21). Glycerate kinase (GLYK) localizes to chloroplasts and functions in photorespiration under light conditions. Under shade conditions, inactivated PHYs lead to alternative transcription initiation and produce cytoplasmic GLYK due to the lack of a cTP for cytoplasmic photorespiratory bypass (21). Consistent with this study, our data show that light-dependent alternative promoter usage makes the *NCP* gene produce NCP-L, which contains a cTP, under high light conditions and the NCP-S isoform without a cTP under low light/dark conditions. However, the impact of this mechanism on NCP is different during photomorphogenesis. Our data indicate that NCP-S, produced without the cTP due to alternative promoter usage during low light/dark, is nonfunctional and degraded by the 26S proteasome-dependent pathway. Organelle-targeted proteins are tightly regulated by the protein import pathway and subjected to protein degradation via the ubiquitin-proteasome system (UPS) if they are mislocalized or misfolded in a eukaryotic cell (27, 28). Therefore, our data suggest that NCP-S, which is produced without a cTP due to alternative promoter usage, is degraded in the cytoplasm. We therefore propose a molecular mechanism in which alternative promoter usage is linked to the quality control of protein localization, ensuring that proteins are correctly targeted to their functional locations and using protein degradation to prevent mistargeting.

Mechanisms for alternative transcription initiation have been proposed at different levels (24, 26). First, transcription factor binding to specific enhancers in different regions can activate alternative transcription (29). Second, structural changes in chromatin via histone modifications or DNA methylation can alter the local architecture and facilitate open chromatin for transcription initiation (30, 31). However, the precise mechanism by which PHYs regulate alternative promoter selection remains elusive. We found that light-dependent alternative transcription initiation in *NCP* is dependent on PIFs, as demonstrated in the *pif1*, *pif3*, and *pifq* mutant backgrounds. Notably, a potential PIF-binding E-box (CANNTG) exists in exon 2, close to TSS2, suggesting that PIF proteins directly bind to and prevent transcription initiation from TSS1. This binding event may inhibit transcription initiation from TSS1 while promoting initiation from TSS2. However, our attempts to identify evidence for direct binding of PIF proteins to the *NCP* gene via chromatin immunoprecipitation (ChIP)-sequencing did not yield conclusive results from the PIF1, PIF3, PIF4, or PIF5 datasets (32–35). It is likely that the PIF-dependent regulation of alternative transcription initiation in the *NCP* gene may involve other transcription factors or chromatin remodelers.

Nucleus-chloroplast dual localization has been previously reported with proteins such as PAP5/HMR, PAP8/pTAC6, and Whirly1 (WHY1) (36–38). Although the mechanism is still unclear, stromules have been proposed as passageways to transport a number of molecules to nuclei (23, 39). Stromules are long, thin, tubular extensions from plastids that provide a potential pathway for retrotranslocated proteins (23, 40, 41). Our data show that the NCP-L isoform appears in the nucleus when the nucleus is surrounded by chloroplasts and when stromules are observed in these chloroplasts. We did not observe stromules when we expressed cTP-YFP in tobacco cells, suggesting that chloroplast-localized NCP may contain an unknown molecular feature necessary to induce stromules. Reactive oxygen species (ROS) increase the frequency of stromule formation in chloroplasts (23, 40). Chloroplasts are a major source of ROS production via the photosynthetic electron transport chain (42, 43). It is possible that chloroplast biogenesis during photomorphogenesis generates ROS as a signal to induce stromules. Notably, dual-targeted proteins such as PAP5/HMR, PAP8/pTAC6, RCB, and NCP, which are essential for chloroplast biogenesis, have been shown to impact PHY signaling by modulating phyB photobody formation in the nucleus (11). The biological significance of chloroplast-to-nucleus retrotranslocation and PHY signaling remains an intriguing question. We propose that during the dark-to-light deetiolation process, when the PEP complex is assembled, chloroplasts transmit a signal to the nucleus via stromules. This signal could facilitate phytochrome signaling by promoting the formation of large photobodies as part of photomorphogenesis. Further research is needed to elucidate the precise molecular mechanisms underlying these processes and their biological significance.

## Materials and methods

### Plant materials and growth conditions

The *ncp-10* (Col-0 background) mutant has been described previously (19). The Arabidopsis mutants *pif1-2* (44), *pif3-1* (45), and *pifq* (46), all in the Col-0 background, were used for the 5’ RACE experiment. Additional Arabidopsis transgenic lines generated in this study are described below. Seeds were surface-sterilized and plated on 1/2X Murashige and Skoog (MS) media with Gamborg’s vitamins (MSP06, Caisson Laboratories, North Logan, UT), 0.5 mM MES pH 5.7, and 0.8% agar (w/v) (A037, Caisson Laboratories). Seeds were stratified in the dark at 4 °C for 4 d. Seedlings were grown at 21 °C in an LED chamber (Percival Scientific, Perry, IA) under the indicated light conditions. For true-dark treatment, stratified seeds were pre-illuminated with far-red light (10 μmol·m^−2^·s^−1^) for 3 h before transferring to darkness. Fluence rates of light were measured using a Li-180 Spectrometer (LI-COR Biosciences., Lincoln, NE).

### Plasmid construction and generation of transgenic plants

To generate the *NCPpro*:*NCP*-*HA*-*His* fusion construct, 3×hemagglutinin (HA) and 6×histidine (His) coding sequences were fused in frame to the 3’ end of the *NCP* genomic sequence driven by its own promoter, consisting of a ∼4.9-kb sequences. The fusion construct was subcloned into the EcoRI and BamHI sites of the *pJHA212G* vector (47) using HiFi DNA assembly (New England Biolabs, Ipswich, MA). The second ATG of the *NCP* coding sequence in the *pJHA212G*-*NCPpro*:*NCP*-*HA*-*His* was mutated to TTG to block the translation of the NCP-S isoform, resulting in *pJHA212G*-*NCPpro:NCP*_2nd ATGm_-*HA*-*His*. Full-size *NCP long* (*L*) and *short* (*S*) coding sequences were fused in-frame to the 5’ end of an *HA-His*-coding sequence, and the fusion protein was overexpressed using the *UBQ10* promoter to generate the *UBQ10*-*NCP*-*L*-*HA*-*His* and *UBQ10*-*NCP*-*S*-*HA*-*His* fusion constructs. Transgenic lines expressing the indicated *NCP*-*HA*-*His* constructs were generated by transforming *ncp-10* plants with *Agrobacterium tumefaciens* strain GV3101 harboring the indicated constructs (48). Multiple independent lines from the T1 generation were selected on MS medium containing 100 μg/ml gentamycin and 2% sucrose. Plants with *ncp-10* mutations were selected in the T1 generation. T2 lines with a single-locus transgene insertion were selected based on a 3:1 segregation ratio for gentamycin resistance. T3 generation plants homozygous for the transgene were used for further experiments. For the *UBQ10*-*NCP*-*S*-*HA*-*His* transgenic lines, plants homozygous for the transgene and heterozygous for the *ncp-10* mutation in the T3 generation were used for subsequent experiments. Primers used for genotyping assays are listed in Supplemental Table 2.

The *NCP*-*L*-*CFP*-*FLAG* and *NCP*-*S*-*CFP*-*FLAG* constructs used for the transient expression assay in tobacco and Arabidopsis protoplast were generated by amplifying the NCP-L and NCP-S coding sequences, and each fragment was fused in-frame to the 5’ end of *CFP*-*FLAG*. The fusion constructs were subcloned into the *pJHA212G* binary vector under the control of the *UBQ10* promoter. All the primers used for plasmid construction are listed in Supplemental Table 1.

### 5’ RACE analysis

One microgram of total RNA was used for first-strand cDNA synthesis for 5’ rapid amplification of cDNA ends (5’ RACE) analysis, which was performed using a SMARTer RACE 5’/3’ Kit (Clontech Laboratories, Mountain View, CA) according to the manufacturer’s instructions. The obtained cDNA was subjected to 5’ RACE-PCR using SeqAmp DNA Polymerase (Takara Bio USA). Primers used for 5’ RACE-PCR and the number of PCR cycles are shown in Supplemental Table 3. The 5’ RACE-PCR products were separated by electrophoresis on a 1% agarose gel, and the DNA contained therein was excised and extracted using a Zymoclean Gel DNA Recovery Kit (Zymo Research, Irvine, CA). Each transcript extracted was then cloned into a pRACE cloning vector (Clontech Laboratories) with an In-Fusion Cloning Kit (Clontech Laboratories). Plasmid DNA was purified using a ZR Plasmid Miniprep-Classic Kit (Zymo Research) and sequenced by the University of Utah DNA sequencing core. At least 3 independent 5’ RACE clones for each reaction were sequenced to define the transcription start sites.

### Transient expression in Arabidopsis protoplasts

To isolate protoplasts from wild-type plants, Col-0 seedlings were grown on 1/2 X MS plates for 2 weeks, and the whole seedlings were pre-incubated in 0.4 M mannitol solution for 1 h. Overnight incubation of the wild-type seedlings was performed in the enzyme solution (20 mM MES pH 5.7, 1.5% (w/v) cellulase R10, 0.4% (w/v) macerozyme R10, 20 mM KCl, 10 mM CaCl2, and 0.1% BSA). High-purity plasmid DNA was prepared using PureLink HiPure Plasmid Midiprep Kit (Thermo Fisher Scientific). Arabidopsis mesophyll protoplast transfection was performed by a polyethylene glycol (PEG)-calcium transfection method (49). The transfection reaction was incubated in white light (100 μmol m^−2^ s^−1^) at room temperature for 24 hours. Protoplasts were mounted, and the subcellular distributions were visualized using a Zeiss Axio Observer 7 inverted microscope with a 40X magnification (Carl Zeiss, Jena, Germany). Chlorophyll autofluorescence was monitored using excitation from a 555/30 nm green LED light with filter set 121. CFP was monitored using 423/44 nm excitation from a violet LED light with filter set 47.

### Transient expression in *N. benthamiana*

*Agrobacterium tumefaciens* strain GV3101 harboring each construct (i.e., *UBQ10*-*NCP*-*L*-*CFP*-*FLAG* and *UBQ10*-*NCP*-*S*-*CFP*-*FLAG*) was grown overnight in 5 ml of LB media and pelleted by centrifugation at 3,000 × g. The bacterial pellet was resuspended in 2.5 ml of infiltration buffer (10 mM MES pH 5.7, 10 mM MgCl_2_) with 200 μM acetosyringone. The bacterial suspension was diluted to an OD_600_ of 1.0 with infiltration buffer and infiltrated into the abaxial side of leaves from 3-week-old tobacco plants. Around 50 h after infiltration, leaf punches were mounted in 1X PBS and imaged on a fluorescence microscope with 40X magnification. DAPI was monitored using 385/30 nm excitation from a UV LED light with filter set 49.

### Hypocotyl measurements and imaging of greening phenotype

For hypocotyl length measurements, 4-d-old seedlings grown under 50 μmol m^−2^ s^−1^ red light were scanned using a Brother MFC-L2750DW scanner, and hypocotyls were measured using ImageJ software (https://imagej.nih.gov/ij/). Box-and-whisker plots of hypocotyl measurements were drawn using Prism 10 software (GraphPad software, Inc., La Jolla, CA). Images of representative seedlings were captured using an AmScope 64-LED stereomicroscope (AmScope, Irvine, CA) and processed using Adobe Photoshop CC (Adobe Systems, Mountain View, CA). For imaging of the greening phenotype used in Figs. 4A and 5A, representative seedlings in the 20 μmol m^−2^ s^−1^ red light condition (R20) were imaged using an AmScope 64-LED stereomicroscope (AmScope).

### RNA extraction and quantitative real-time PCR

Preparation of RNA samples from seedlings of the indicated genotypes and growth conditions was performed using a Quick-RNA MiniPrep Kit with on-column DNase I treatment (Zymo Research). cDNA synthesis was performed with 1.5 µg of total RNA using a Superscript IV First-Strand cDNA Synthesis Kit (Thermo Fisher Scientific) according to the manufacturer’s protocol.

Oligo(dT) primers were used for the analysis of transgene expression in the nucleus, and a mixture of plastidial-gene-specific primers was used to analyze plastidial genes. qRT-PCR reactions were performed in 96-well plates on a LightCycler 96 System (Roche, Basel, Switzerland) using FastStart Essential DNA Green Master (Roche) in a volume of 10 μL. The *PP2A* gene was included as an internal control in the PCR reactions to normalize for variations in the amounts of cDNA used. Primers for QRT-PCR and cDNA synthesis are listed in Supplemental Table 2.

### Protein extraction and immunoblot analysis

Total protein was extracted from Arabidopsis seedlings grown under the indicated conditions. Plant tissues were ground in liquid nitrogen and resuspended in extraction buffer (100 mM Tris-HCl pH 7.5, 100 mM NaCl, 1% SDS, 5 mM EDTA pH 8.0, 20 mM DTT, 40 μM MG132, 40 μM MG115, and EDTA-free protease inhibitor cocktail) at room temperature, then centrifuged at 20,000 × g for 10 min. The supernatant was mixed with the same volume of 2X SDS loading buffer and boiled for 10 min. Protein extracts were separated on an SDS-PAGE mini-gel and transferred onto a polyvinylidene difluoride (PVDF) membrane. The membrane was blocked in 5% non-fat milk in 1X TBST and detected with mouse monoclonal anti-HA (H9658, Sigma-Aldrich, St. Louis, MO) and anti-actin (A0480, Sigma-Aldrich) antibodies. An anti-mouse (1706516, Bio-Rad, Hercules, CA) secondary antibody was used at a 1:5000 dilution. The signals were detected with a chemiluminescence reaction using SuperSignal West Dura Extended Duration Chemiluminescent Substrate (Thermo Fisher Scientific).

## Acknowledgements

This work was supported by start-up funds from the University of Utah to C. Y. Y. We thank Yejin Park for imaging processes used in Figure 4C and Figure 5B. We thank Elise Pasoreck and Heejin Yoo for valuable comments and suggestions on the manuscript.

## Author contributions

J.-H.L. and C.Y. designed research; J.-H.L., T.M.D, and S.S. performed research; J.-H.L and C.Y. analyzed data; and J.-H.L. and C.Y. wrote the paper.

