## Supplemental Figures and Tables for "Mechanism of nucleus-chloroplast communication by alternative promoter usage and stromules to establish photomorphogenesis in Arabidopsis"

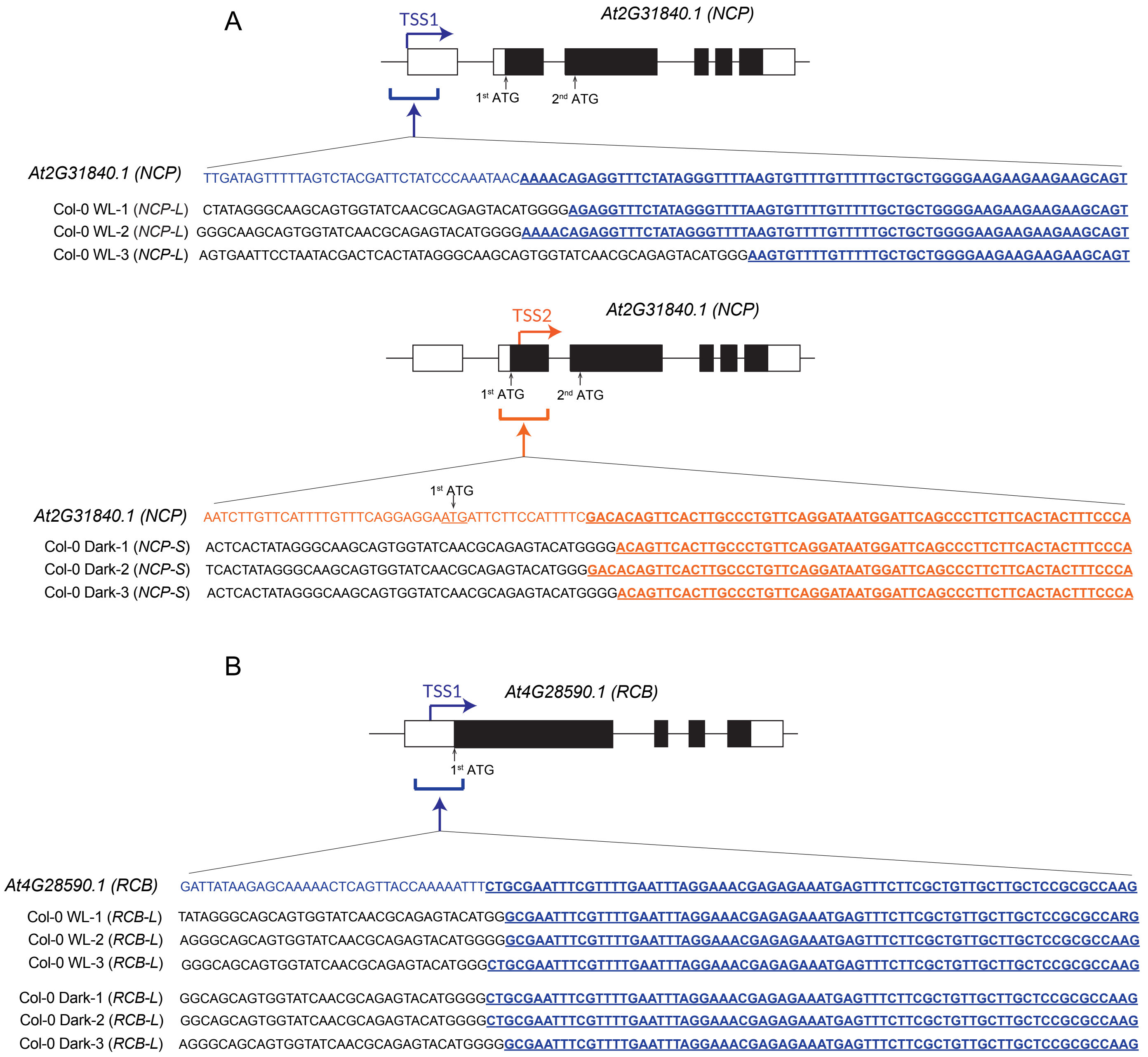


**Supplemental Figure 1. Representative Sanger sequencing results for *NCP* and *RCB* 5' RACE cDNA clones.**

(A) Identification of *NCP* 5' cDNA ends with different transcription start sites (TSSs). Each *NCP* transcript amplified from 5' RACE-PCR was extracted and subcloned into a linearized pRACE vector (Clontech). Three independent *NCP* 5' RACE clones for each reaction were sequenced. Blue and red brackets indicate regions showing sequence aligned. The underlined blue characters indicate matched DNA sequences from *NCP-L* transcript with TSS1 and red underlined characters denote aligned sequences from *NCP-S* transcript with TSS2. The *NCP-S* transcript started with TSS2 in the region after 1st ATG in the coding sequence. The black DNA sequences for each reaction are resulted from SMARTer 5' RACE cDNA synthesis (Clontech).

(B) Identification of *RCB* 5' cDNA end. Preparation of 5' RACE clones were described as in (A). Three independent 5' RACE clones for each reaction were sequenced. Blue bracket indicates region sequence aligned. The underlined blue characters denote matched sequences from *RCB-L* transcript with annotated *RCB* sequence. The TSS1 of *RCB* was verified in downstream of the annotated TSS. The black DNA sequences are resulted from SMARTer 5' RACE cDNA synthesis.

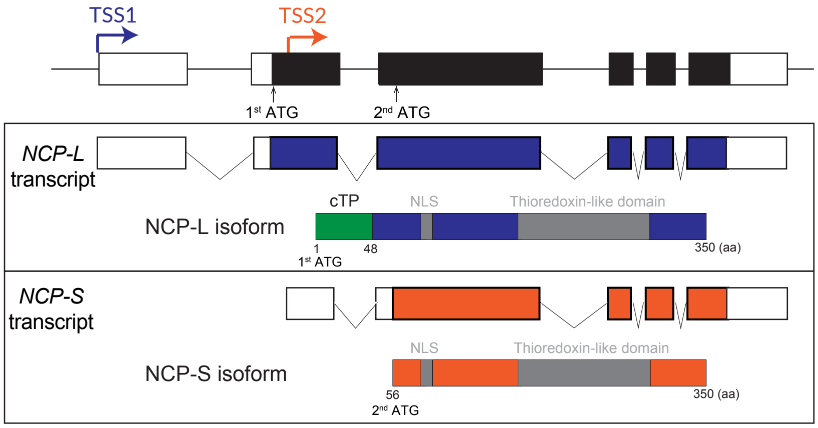


**Supplemental Figure 2. Schematic illustration of the predicted domain structure of NCP-L and NCP-S isoforms.** cTP, chloroplast-targeting transit peptide; NLS, putative nuclear localization signal.

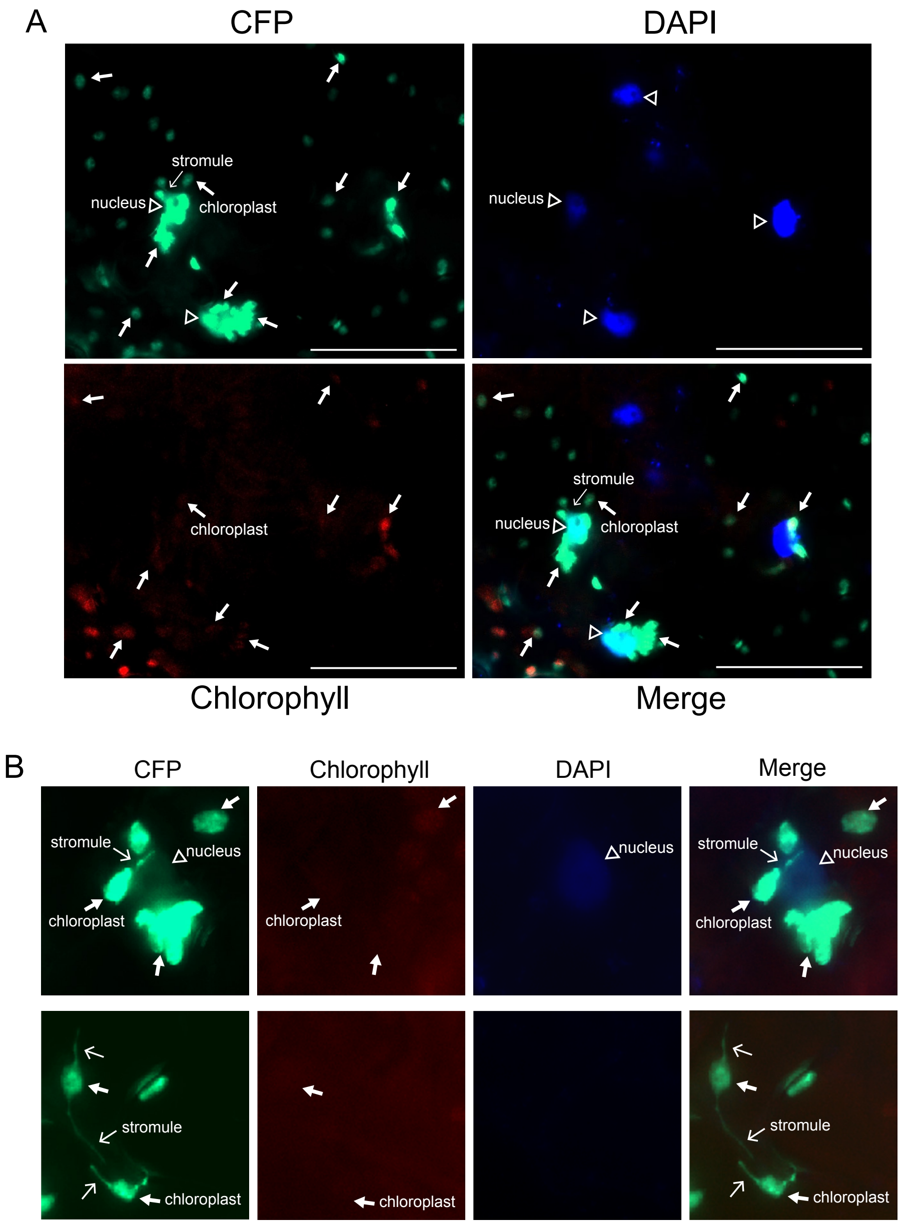


**Supplemental Figure 3. Localization patterns of NCP-L isoform in *Nicotiana benthamiana* leaves.**

(A) Uncropped fluorescence microscopy images of NCP-L-CFP. The *UBQ10-NCP-L-CFP-FLAG* fusion construct was expressed transiently in tobacco leaves. Fluorescence microscopy images of epidermal cells were visualized. NCP-L-CFP signal was mostly detected in chloroplasts (thick arrows) and often observed in the nucleus (empty arrowheads) when they are surrounded by chloroplasts. Thin arrow denotes stromule. DAPI was used for staining nuclei. Chlorophyll indicates autofluorescence. Scale bars, 50 $\mu$m.

(B) Expression patterns of NCP-L isoform in stromules. The stromule expressed NCP-L-CFP was conneted with nuclues (upper panel). NCP-L-CFP was expressed in chloroplast visualizing stromules on the both side of chloroplast (lower panel).

**Supplemental Table 1.** Primers used in construct preparation.

| Construct | Primer | Sequences (5’- 3’) | Usage |
|---|---|---|---|
| *UBQ10-NCP-L-CFP-FLAG* | NCP longer CDS_fwd | CAGCGAGCTCGGTACCCGGGATGATTCTTCCATTTTCGAC | Subcloning |
| *UBQ10-NCP-L*(*S*)*-CFP-FLAG* | NCP CDS_rev | CTTTACTCATATAATTCACACTTACATCGAC | Subcloning |
| *UBQ10-NCP-L*(*S*)*-CFP-FLAG* | sCFP3A 3FLAG_fwd | TGTGAATTATATGAGTAAAGGAGAAGAAC | Subcloning |
| *UBQ10-NCP-L*(*S*)*-CFP-FLAG* | sCFP3A 3FLAG_rev | GAAAGCTCTGCATGCCTGCATCACTTGTCATCGTCATC | Subcloning |
| *UBQ10-NCP-S-CFP-FLAG* | NCP shorter CDS_fwd | CAGCGAGCTCGGTACCCGGGATGAGGTCGAGGAGAAATG | Subcloning |
| *NCPpro:NCP-HA-His* | gNCP_fwd | ACAGCTATGACATGATTACGTTATCACATCCATTCACATTTG | Subcloning |
| *NCPpro:NCP-HA-His* | gNCP_rev | CCCGGGTACCATTCACACTTACATCGAC | Subcloning |
| *NCPpro:NCP-HA-His* | HA-His_fwd | AAGTGTGAATGGTACCCGGGGATCCTCTAG | Subcloning |
| *NCPpro:NCP-HA-His* | HA-His_rev | CCTGCAGGTCGACTCTAGAGTCAGTGATGGTGATGGTGATG | Subcloning |
| *NCPpro:NCP 2nd ATGm-HA-His* | NCP 2nd ATGm _fwd | AAGTGGAATGTTTTGAGGTCGAGGAGA | Subcloning |
| *NCPpro:NCP 2nd ATGm-HA-His* | NCP 2nd ATGm _rev | TCTCCTCGACCTCAAAACATTCCACTT | Subcloning |
| *UBQ10-NCP-L-HA-His* | NCP longer CDS_fwd | CAGCGAGCTCGGTACCCGGGATGATTCTTCCATTTTCGAC | Subcloning |
| *UBQ10-NCP-L*(*S*)*-HA-His* | NCP CDS_rev | CCCGGGTACCATTCACACTTACATCGAC | Subcloning |
| *UBQ10-NCP-L*(*S*)*-HA-His* | HA-His_fwd | AAGTGTGAATGGTACCCGGGGATCCTCTAG | Subcloning |
| *UBQ10-NCP-L*(*S*)*-HA-His* | HA-His_rev | GAAAGCTCTGCATGCCTGCATCAGTGATGGTGATGGTGATG | Subcloning |
| *UBQ10-NCP-S-HA-His* | NCP shorter CDS_fwd | CAGCGAGCTCGGTACCCGGGATGAGGTCGAGGAGAAATG | Subcloning |

The PCR primers were designed using NEBuilder software (version 2.10.1, https://nebuilder.neb.com/#!/) so that they have calculated melting temperatures in the range of 55−65$^{o}$C. fwd, forward primer; rev, reverse primer.

**Supplemental Table 2.** Primers used in genotyping PCR, cDNA synthesis, and RT-qPCR.

| Primer | Sequences (5'-3') | Usage |
|---|---|---|
| *ncp-10*_LP | AAGGGAGAAGAGAGCACGTT | Genotyping |
| *ncp-10*_RP | GTAGAGAGACGGGAATGGAGG | Genotyping |
| *ncp-10*_LB | ATAATAACGCTGCGGACATCTACATTTT | Genotyping |
| *NCP-L(S)-HA-His* transgene_fwd | GTAGAGAGACGGGAATGGAGG | Genotyping |
| *NCP-L(S)-HA-His* transgene_rev | GAAAGCTCTGCATGCCTGCATCAGTGATGGTGATGGTGATG | Genotyping |
| psbA_cDNA | TAGATGGAGCCTCAACAGCAGCTA | cDNA synthesis |
| rbcL_cDNA | CTTCACAAGCAGCAGCTAGTTCAGG | cDNA synthesis |
| psbA_fwd | ACATTTCTTCTTAGCGGCTT | RT-qPCR |
| psbA_rev | CGTCCTTGACTATCAACTACTGA | RT-qPCR |
| rbcL_fwd | GGAGATGATTCTGTACTACAAT | RT-qPCR |
| rbcL_rev | GTCCCTCATTACGAGCTTGTAC | RT-qPCR |
| *NCP-L(S)-HA-His* transgene_fwd | CTTGTTTGAATGGTGTCGTGAAT | RT-qPCR |
| *NCP-L(S)-HA-His* transgene_rev | GTTGGTGTAGGTGTTGGAGTAG | RT-qPCR |
| PP2A_fwd | TATCGGATGACGATTCTTCGTGCAG | RT-qPCR |
| PP2A_rev | GCTTGGTCGACTATCGGAATGAGAG | RT-qPCR |

The PCR primers were designed using Primer3 software (version 4.1.0, https://primer3.ut.ee/) in a way that they have calculated melting temperatures in a range of 50-60oC. fwd, forward primer; rev, reverse primer.

**Supplemental Table 3.** Primers used for 5' RACE-PCR and the number of PCR cycles.

| Primer | Sequences (5'-3') | PCR cycles |
|---|---|---|
| Universal Primer A Mix (UPM) | CTAATACGACTCACTATAGGGCAAGCAGTGGTATCAACGCAGAGT | 30 cycles |
| NCP 5' RACE_GSP | GATTACGCCAAGCTTACTTGAATCGCCTTCTCTAGTTCTTCCC | 30 cycles (with UPM) |
| Universal Primer Short (UPS) | CTAATACGACTCACTATAGGGC | 25 cycles |
| NCP 5' RACE_NGSP | GATTACGCCAAGCTTGTCTTCCTCCATTCCCGTCTCTCTACTG | 25 cycles (with UPS) |
| RCB 5' RACE_GSP | GATTACGCCAAGCTTCTCCCTGTACAGAATCCGGT | 30 cycles (with UPM) |
| RCB 5' RACE_NGSP | GATTACGCCAAGCTTGATTTACGCGGGACCTTTGT | 20 cycles (with UPS) |
| PP2A_fwd | TGCCCCAGATGTGCTAAAGA | 25 cycles |
| PP2A_rev | GCTGCTATCCGAACTTCTGC | 25 cycles |
